# Deep learning inference of universal dormancy pseudotime reveals the cellular targets of anti-cancer therapies

**DOI:** 10.64898/2025.12.14.692655

**Authors:** Haley M. Beckmann, Michelle Tong, Glenn Chang, Adi Steif

**Affiliations:** Department of Basic and Translational Research, BC Cancer Research Institute, Vancouver, BC, Canada V5Z 1L3; Department of Medical Genetics, University of British Columbia, Vancouver, BC, Canada V6T 1Z4; Canada’s Michael Smith Genome Sciences Centre, Vancouver, BC, Canada V5Z 4S6

**Keywords:** dormancy pseudotime, quiescence, senescence, single cell transcriptomics, deep learning, immune checkpoint inhibition, platinum chemotherapy, breast cancer, ovarian cancer

## Abstract

Controlled exit from and re-entry into the cell cycle is essential for multi-cellular life, while aberrant quiescent and senescent cell states have been implicated in age-related diseases and cancer treatment evasion. Recent molecular and imaging studies suggest non-cycling cellular states exist along a continuum of deepening dormancy, whereby the probability of cell cycle re-entry decreases with distance from the restriction point. We trained a probabilistic deep-learning model that enables mapping of heterogeneous single cell transcriptomic datasets into an interpretable latent space that encodes a common “dormancy pseudotime”. We demonstrate that our model enables robust inference of active cell cycle states, and validate in diverse biological contexts that the inferred location along dormancy pseudotime represents a continuum from quiescence to durably arrested states. Applying dormancy pseudotime inference to pre- and post-treatment time points from patients undergoing anti-cancer treatment, we uncover new insights into the distinct tumour cell dormancy states targeted by immune checkpoint inhibitors and platinum-taxane chemotherapy. Given the ubiquity of single cell transcriptomics, we anticipate that dormancy pseudotime analysis will be widely applied to shed new light on the complex interplay between cycling and non-cycling cellular states in health and disease.

## Introduction

The regulation of cell cycle dynamics including progression through, exit from and re-entry into the cell cycle is essential for development and differentiation^1, 2^, tissue homeostasis^3^, wound repair^3^, immune function^3^, and healthy aging^2^. Cell cycle dysregulation is a central hallmark of cancer^4^, while the presence of quiescent tumour cells has been associated with therapeutic resistance and disease recurrence^5–7^. With the arrival of high-throughput single cell RNA sequencing (scRNA-seq), numerous methods were developed to detect cells in the active phases of the cell cycle^8–10^ (G1, S and G2/M), with the primary aim of removing unwanted sources of variation that can confound cell clustering. More recently, several tools were introduced that place actively proliferating cells along continuous circular trajectories, and infer “cell cycle pseudotime”^11–16^. These methods enable deeper characterization of cell-cycle-associated genes and their dynamics over the cell cycle, and work effectively when applied to datasets comprised primarily of actively proliferating cells from a single cell type. However, these tools assume the presence of a prominent cyclical manifold within the data. This limits their utility in the context of transcriptomic datasets derived from primary tissues, which feature numerous cell types and are often composed primarily of non-cycling cells.

At the restriction point in G1, cells must commit to active growth and cell cycle re-entry, or exit to a state outside of the cell cycle termed G0^3^. This decision can be developmentally programmed or triggered by intra- and extra-cellular conditions, and may be reversible or permanent^3^. Quiescence, a state of non-proliferation that is reversible in response to an appropriate stimulus, is actively maintained in a variety of cell types^3^. This includes, for example, tissue-resident adult stem cells, which provide an essential reservoir for tissue repair and regeneration^3^, as well as naive and memory lymphocytes, which while inactive are poised to rapidly proliferate in response to antigen stimulation^17^. Irreversible cell cycle exit, on the other hand, is characteristic of terminal differentiation and senescence^3^. While canonically considered distinct states, recent studies based on cell imaging and protein expression suggest that quiescence and senescence exist along a G0 continuum, whereby a cell’s likelihood of cell cycle re-entry decreases with distance from the restriction point^18, 19^. While cells in quiescent and senescent states can feature pronounced differences in morphology and metabolic activity, both are accompanied by a downregulation of cell cycle associated genes and activation of regulatory pathways including p53-p21, p16-RB-E2F, and p27^2,3,19,20^.

We hypothesize that despite heterogeneous functional states and conditions triggering cell cycle exit, the transcriptomic profiles of G0 cells from diverse cell types and tissue contexts can be projected onto a common trajectory capturing depth of dormancy, which we term “dormancy pseudotime”. We introduce Ouroboros, a computational model and analysis package for single cell RNA sequencing (scRNA-seq) data that leverages a variational autoencoder (VAE) with learned reference embedding to place both non-cycling and actively proliferating cell populations along continuous dormancy and cell cycle trajectories. Recently introduced computational approaches discriminate non-cycling G0 cells from active cell cycle states in scRNA-seq data and characterize G0 populations^21, 22^. However, our model is first to infer cellular location along a common dormancy pseudotime. In determining the relative position of cells within a shared latent space, dormancy pseudotime analysis reveals subtle shifts in non-cycling cell state distributions between populations, and permits quantitative downstream analyses of associated gene expression changes. On an independent FUCCI-labeled dataset, our model shows robust performance in classifying active cell cycle phases and cell cycle pseudotime. Applying dormancy pseudotime analysis to scRNA-seq data from cells exposed to senescence-inducing stimuli, we validate the association between deepening dormancy pseudotime and the expression of senescent gene signatures. We demonstrate that samples comprising heterogeneous cell types, including a well-characterized mouse pancreatic endocrinogenesis dataset, are well-integrated within the model’s latent space. Finally, we apply dormancy pseudotime inference to scRNA-seq data from two anti-cancer drug studies with pre- and post-treatment timepoints, revealing unexpected shifts in tumour cell dormancy distributions in response to immunotherapy and chemotherapy, as well as an association between post-treatment cellular dormancy states and patient outcomes.

## Results

### Ouroboros latent space encodes cell cycle dynamics independent of cell type and technology

We developed Ouroboros, a universal cell cycle and dormancy state embedding, by training a variational autoencoder (VAE) to project cells onto the surface of a spherical latent space^23, 24^. The probabilistic nature of VAEs enables embedding of previously unseen data points into the learned embedding, where their position relative to neighbouring training data points can be used to infer a discrete phase label, cell cycle pseudotime, and dormancy pseudotime (**Figure 1A**).

**Figure 1.**
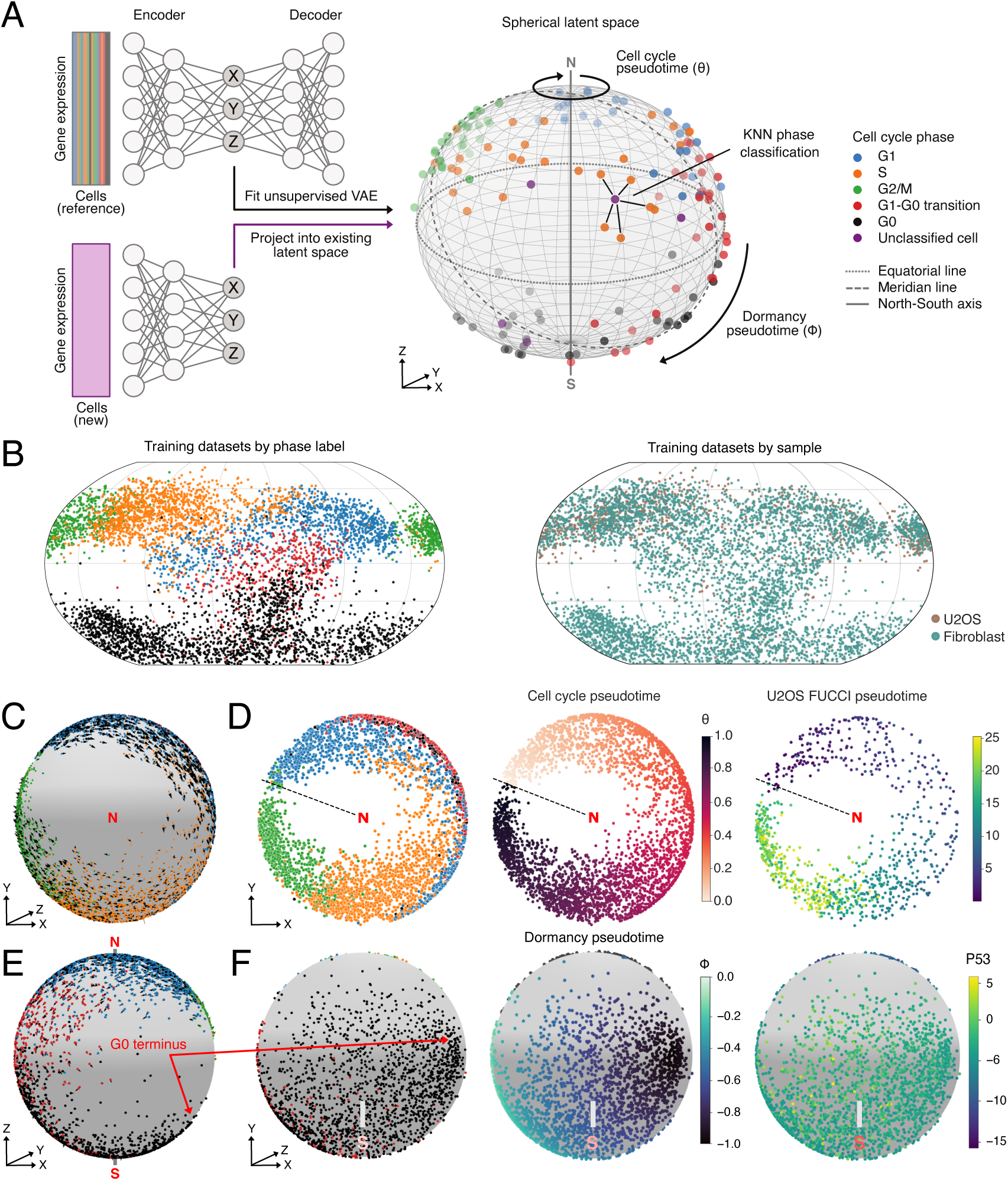
Ouroboros is a trained VAE with interpretable spherical latent space that captures cell cycle and dormancy states. (**A**) Model schematic. The unsupervised VAE is trained on a scRNA-seq data with known cell cycle phase. Using the trained encoder, new datasets are projected into the latent space, where they are assigned a KNN phase label and either a cell cycle or dormancy pseudotime based on their position relative to the training data. (**B**) The U2OS (n = 1021 cells) and primary fibroblast (n = 4677 cells) training datasets in VAE latent space, coloured by phase label (left), and sample (right). (**C**) RNA velocity around the North-South axis for cycling cells in the northern hemisphere of the sphere. (**D**) Cells in the training dataset are projected onto the equatorial plane and assigned a cell cycle pseudotime (*θ*, range 0 to 1), which is strongly correlated with FUCCI pseudotime. Dotted line represents *θ* = 0. (**E**) RNA velocity for non-cycling cells along the meridian line of the sphere. The “G0 terminus” in the embedding is associated with deepest dormancy. (**F**) Cells in the training dataset are assigned a dormancy pseudotime (*ϕ*, range 0 to −1) based on their position between the equatorial plane and “G0 terminus”. P53 shows elevated expression in the southern hemisphere of the sphere.

To train the VAE, we selected two well-characterized datasets that represent the cell cycle across different biological and technological contexts. The first is a SMART-seq2 dataset derived from the U2OS human osteosarcoma cell line with orthogonal fluorescent ubiquitination-based cell cycle indicator (FUCCI) labels and pseudotime, as well as a previously described non-cycling sub-population^25^ (n = 1021). The second is a 10x Genomics Chromium dataset comprised of cultured primary human fetal lung fibroblasts without orthogonal cell cycle phase labels, which was also previously reported to feature a non-proliferative cell subset^16^ (n = 4677). We assigned discrete phase labels based on marker expression and RNA velocity, identifying the non-proliferative (G0) cell subsets and an intermediate G1-G0 transition state. The training datasets integrated well in the VAE latent space, with overlapping phase labels in distinct regions of the sphere (**Figure 1B**).

Within the trained embedding, cells in G1, S, and G2/M phases formed a circle around the “northern hemisphere” of the sphere, and showed RNA velocity consistent with transit through the cell cycle (**Figure 1C**, **Figure S1**). RNA velocity magnitude became very small and had no discernible direction in late G2/M, which is consistent with the pause in transcription known to occur in this phase^26, 27^ (**Figure S2**). A clear gap was observed between G2/M and G1, corresponding to telophase and cytokinesis. This is followed by a resumption of directional RNA velocity in G1 (**Figure S2**).

We defined cell cycle pseudotime (*θ* ∈ [0, 1]) based on the polar angle for cells in the northern hemisphere of the latent space (**Figure 1D**). The start and end point, *θ* = 0 = 1, was aligned with the gap between G2/M and G1, with cell cycle pseudotime increasing clockwise around the equatorial plane. For the U2OS cell line, where orthogonal cell cycle marker gene expression was available, Ouroboros cell cycle pseudotime was strongly correlated with FUCCI pseudotime (**Figure 1D**, **Figure S3**, Pearson’s *ρ* = 0.875, p = 7.96 × 10*^−^*^284^).

In Ouroboros latent space, G1-G0 transition cells bifurcated from the cell cycle below the G1-phase and showed RNA velocity consistent with cell cycle exit, pointing downwards towards the “southern hemisphere” of the sphere (**Figure 1E**,**F**, **Figures S1**, **S4**). Cells in G0 dominated the embedding around the southern pole, and extended beyond this point to a “G0 terminus” that remained well-separated from the active cell cycle (**Figure 1E**,**F**, **Figures S1**, **S4**). The southern hemisphere showed elevated TP53 expression (**Figure 1F**), while well-characterized cell cycle genes^28^ and genes downregulated in quiescence^29^ decreased rapidly along this southern trajectory (**Figure S5**). We therefore hypothesized that the trajectory from the G1 bifurcation point to towards “G0 terminus” represents the latent continuum of deepening dormancy states^18, 19^. We defined dormancy pseudotime (*ϕ* ∈ [−1, 0], **Figure 1F**) based on position along this trajectory.

### Ouroboros outperforms existing tools in cell cycle phase classification and pseudotime inference

To benchmark Ouroboros’ performance in inferring active cell cycle states and cell cycle pseudotime, we projected an independent test dataset with FUCCI marker expression derived from the retinal pigment epithelial cell line RPE1 and sequenced using CEL-Seq2 and scEU-seq^30^ (n = 6070) into the model’s latent space. FUCCI pseudotime was obtained from Battich et al.^30^ To assign ground-truth cell cycle phase labels (G1, S, and G2/M) we applied the same FUCCI marker expression thresholds used by Mahdessian et al.^25^ (**Figure S6**), while non-cycling cells were labelled using the same strategy as the training set, based on RNA velocity and marker expression.

We projected the RPE1 cells onto the pre-trained Ouroboros latent space, inferred cell cycle and dormancy pseudotime, and used a K-nearest-training-set-neighbours strategy to assign discrete phase labels (**Figure 2A**). We then compared Ouroboros’ performance against that of three existing cell cycle inference tools: Scanpy^10^, Tricycle^15^, and DeepCycle^16^. Scanpy offers a popular function for scoring cells using well-established S and G2/M marker genes^28^. It outputs discrete cell cycle phase assignments, but does not output cell cycle pseudotime or distinguish G0 cells. Tricycle and DeepCycle use deep learning embeddings to infer cell cycle pseudotime and assign discrete phase labels. However, neither tool classifies G0 cells or infers dormancy pseudotime. All tools were run with default parameter settings, with the exception of DeepCycle which was run with a lower count threshold for training genes due to the overall low total counts in this dataset.

**Figure 2.**
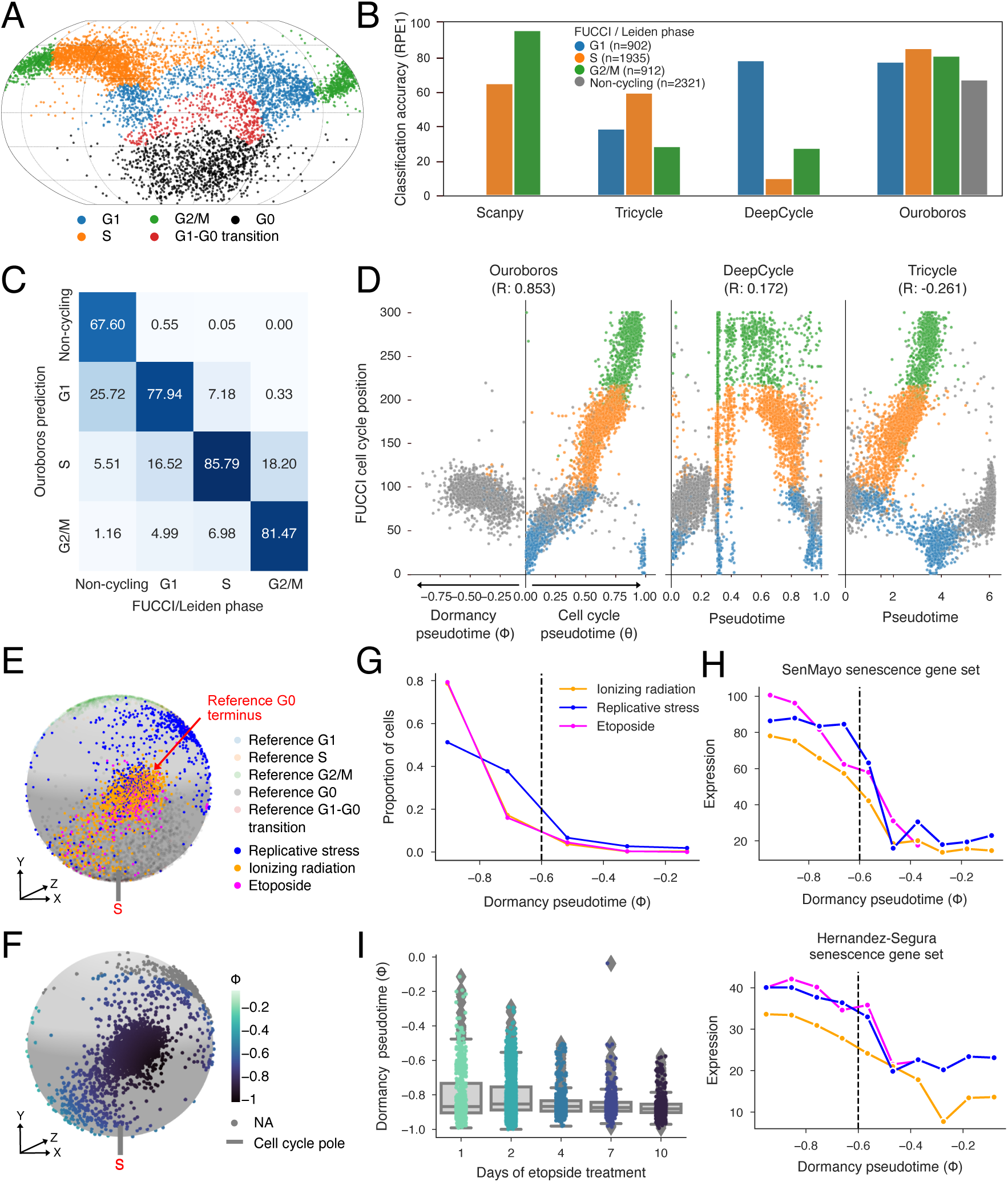
Evaluating Ouroboros using a FUCCI-labelled RPE1 dataset (n=6070 cells) and an induced-senescence dataset (n=7116 cells). (**A**) The RPE1 dataset in Ouroboros latent space, coloured by FUCCI/Leiden phase. (**B**) Benchmarking classification accuracy for Ouroboros and three established methods on the RPE1 dataset. (**C**) Confusion matrix comparing Ouroboros predicted phase labels to ground-truth FUCCI (G1, S, G2/M) and Leiden (non-cycling) labels. Most errors occur between adjacent phases. (**D**) Benchmarking pseudotime inferred by Ouroboros, DeepCycle, and Tricycle relative to FUCCI pseudotime, with points coloured by FUCCI / Leiden labels. R, Spearman’s correlation. (**E**, **F**) Spherical latent space embeddings for the induced-senescence dataset, coloured by treatment condition (top) and dormancy pseudotime (bottom). (**G**, **H**) Proportion of cells (left) and expression of senescence gene sets (right) across dormancy pseudotime bins by treatment condition. Dotted line represents the threshold for deep dormancy. (**I**) A time series of induced senescence via etoposide shows decreasing dormancy pseudotime over 10 days of treatment.

Ouroboros achieved 79% accuracy when classifying discrete cell cycle phases (**Figure 2B**,**C**). We note that the vast majority (74%) of cells “misclassified” by Ouroboros were assigned to a neighbouring phase in the cell cycle trajectory (e.g. G1-S, S-G2/M, **Figure 2C**). This emphasizes the inherent challenge in benchmarking discrete class assignments for a continuous biological process. Scanpy, Tricycle and DeepCycle all had lower discrete phase classification accuracies than Ouroboros (38%, 34% and 28% respectively; **Figure 2B**, **Figure S7**), though their performance improved when non-cycling cells were excluded from the analysis (39%, 47%, and 39% respectively).

Cell cycle pseudotime inferred by Ouroboros was strongly correlated with FUCCI pseudotime for the RPE1 dataset (Pearson’s *ρ* = 0.853, p = 0.00; **Figure 2D**). Tricycle and DeepCycle both showed weak correlations between inferred cell cycle pseudotimes and FUCCI pseudotime (Tricycle: Pearson’s *ρ* = −0.261, p = 0.00; DeepCycle: Pearson’s *ρ* = 0.171, p = 3.18 × 10*^−^*^41^; Figure 2D).

### Induced senescent cells map to Ouroboros deep dormancy pseudotime

To determine whether progression along Ouroboros dormancy pseudotime captures the G0 continuum reported in past live-cell imaging studies, we embedded a dataset profiled using 10x Genomics Chromium from the WI-38 cell line into Ouoroboros latent space^31^ (**Figure 2E**). Wechter et al. induced senescence in this fibroblast cell line via replicative stress (RS), ionizing radiation (IR), or etoposide treatment^31^. As expected, we found that the majority of cells from all three senescence-inducing treatment conditions mapped near the “G0 terminus”, corresponding to the deepest point in Ouroboros dormancy pseuduotime (**Figure 2E**,**F,G**). A time-series experiment with etoposide showed gradual progression towards deeper dormancy pseudotime over 10 days of treatment (**Figure 2I**, **Figure S8**). Finally, we examined the expression of two published gene sets upregulated in senescence (SenMayo^32^ and Hernandez-Segura^33^), both of which increased with deepening dormancy pseudotime (**Figure 2H**). We noted that both the inflection point for the proportion of induced-senescent cells (**Figure 2G**) and the region of highest expression for senescence gene sets, corresponded to the pseudotime range *ϕ* ∈ [−1, −0.6). In subsequent analyses, we refer to cells within this pseudotime range as being in “deep dormancy”.

### Ouroboros recapitulates cycling and non-cycling dynamics in datasets derived from complex tissues

To determine whether Ouroboros latent space captures cell cycle and dormancy states in datasets derived from primary tissues with diverse cell types, we next applied our method to a well-characterized mouse pancreatic endocrinogenesis dataset^34^. A frequently reproduced UMAP projection^35–37^ shows a large cluster of actively cycling ductal progenitor cells at one end of the trajectory, as well as a small cluster of actively cycling endocrine precursor cells further along the differentiation path (**Figure 3A**,**C**). While the UMAP projection captures cell type differences and recapitulates differentiation dynamics, when projected to Ouroboros latent space, all cycling cells were well-integrated with the training dataset in the northern hemisphere of the sphere (**Figure 3B**,**D**). RNA velocity also captured cell cycle progression and exit in this representation, rather than differentiation dynamics (**Figure 3B**,**E**). All but one of the actively cycling cells in this dataset belong to precursor cell types, whereas cells further along the differentiation trajectory predominantly mapped to the G1-G0 transition or the upper G0 regions of the embedding (**Figure 3B**,**E**). We further note that in the Ouroboros embedding, the quiescent cell population also represents a well-integrated mixture of the differentiated cell types, in contrast to the branching pattern observed in the UMAP (**Figure 3A**,**B**). Consistent with expectations, none of the cells in this early embryogenesis dataset mapped to deep dormancy pseudotime (**Figure 3F**). The median dormancy pseudotime for G0 cells in this embryonic dataset was *ϕ* = −0.21, which falls within what we will hereafter refer to as light dormancy (*ϕ* ∈ [−0.4, 0]). We then define mid-dormancy as (*ϕ* ∈ [−0.6, −0.4)). This analysis further demonstrates that the location along dormancy pseudotime in Ouroborus latent space is reflective of cell states across multiple species, cell types, and biological contexts.

**Figure 3.**
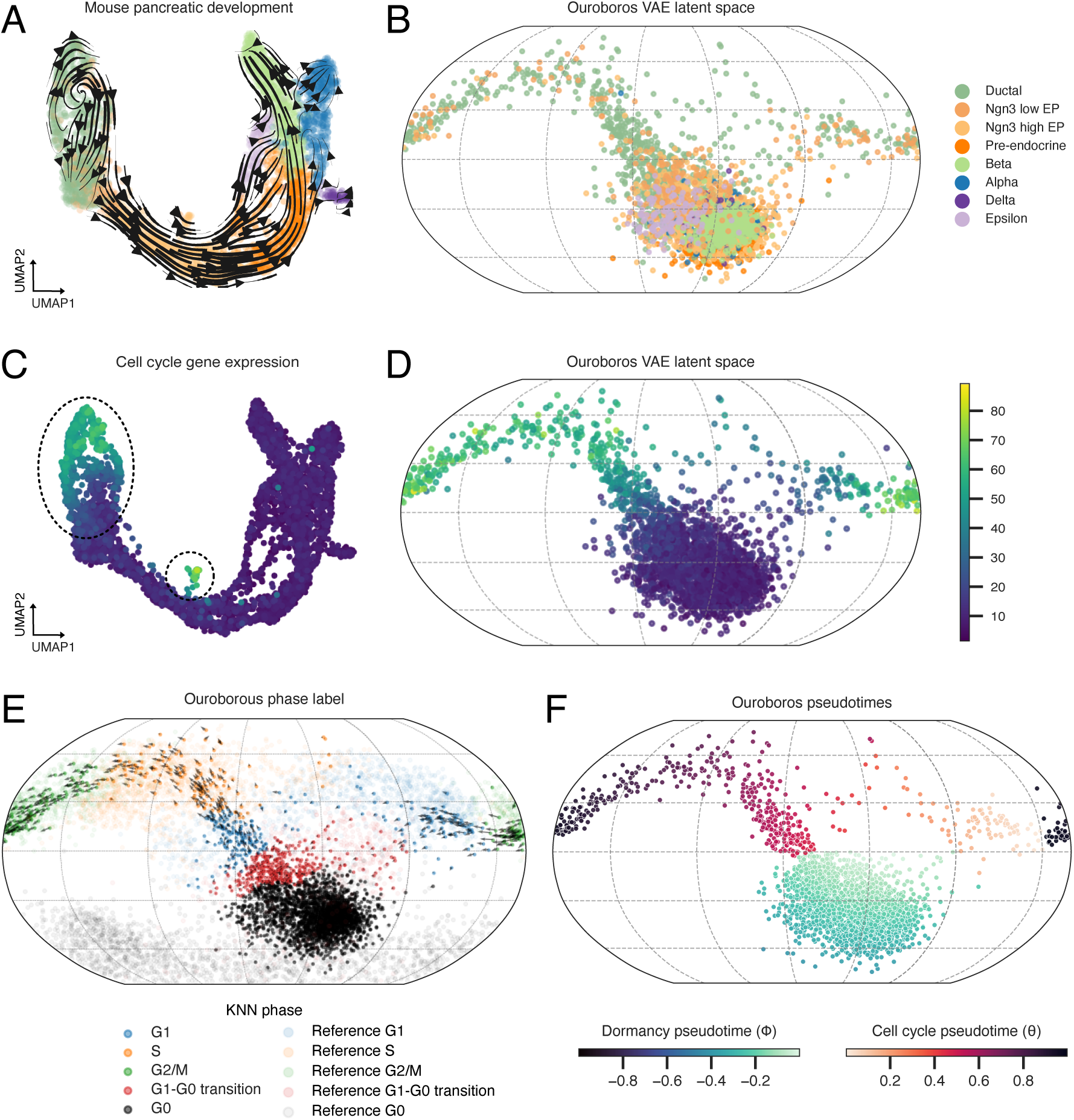
Ouroboros captures cell cycle and dormancy dynamics in a complex mouse pancreatic endocrinogenesis dataset (n=3551 cells). (**A**) UMAP projection and RNA velocity, coloured by cell type. The trajectory is dominated by differentiation dynamics. (**B**) Ouroboros VAE latent space embedding, coloured by cell type. (**C**) UMAP projection, coloured by the expression of cell cycle genes. Two cell subsets in distinct regions of the differentiation trajectory (circled) show high cell cycle gene expression. (**D**) Cycling and non-cycling cells are well-integrated in the northern and southern hemispheres of Ouroboros latent space, respectively. (**E**) RNA velocity in Ouroboros latent space is dominated by cell cycle dynamics in the northern hemisphere, and cell cycle exit towards the southern hemisphere. Pancreatic cells are coloured by KNN phase. Reference cells shown with low opacity. (**F**) Ouroboros pseudotimes in the VAE latent space. Consistent with early embryogenesis, none of the cells in this dataset map to deep dormancy pseudotime.

### Immunotherapy-induced T cell response preferentially targets breast cancer cells in deep dormancy

Next, we sought to determine whether Ouroboros could reveal shifts in cancer cell dormancy pseudotime distributions in response to treatment. We applied our model to primary breast cancer (BC) tumours profiled with 10x Genomics Chromium single cell RNA and T cell receptor sequencing (n = 29 patients, 172,700 cells)^38^. For each patient, a tumour sample was collected before treatment (“pre-treatment” time point) and during treatment with anti-PD1 immune checkpoint inhibitors (ICI) (“on ICI” time point)^38^. Bassez et al. identified 9 patients as having undergone T cell clonotype expansion, hereafter referred to as “responders”, and 20 patients with limited to no T cell expansion, hereafter referred to as “non-responders”^38^. Tumour samples were well-integrated in Ouroboros latent space (**Figure S9**). As expected, we found that responders had a higher proportion of proliferating T cells relative to total T cells both pre- and on-ICI compared to non-responders (Wilcoxon rank-sum test: pre-treatment, U = 161.0, p = 0.00036; on-ICI, U = 153.0, p = 0.0020; **Figure S10**).

Comparing tumour cell dormancy pseudotime distributions, we found that responders had a significantly higher proportion of cancer cells in deep dormancy than non-responders (Wilcoxon rank-sum test: U = 533.0, p = 0.0031; **Figure 4A**,**B**). With treatment, the dormancy pseudotime distribution for responder tumour cells shifted towards lighter dormancy, indicating that T cell expansion is accompanied by a loss of cells in deep dormancy (**Figure 4A**). This shift was not observed for non-responder tumour cells across treatment timepoints (**Figure 4A**).

**Figure 4.**
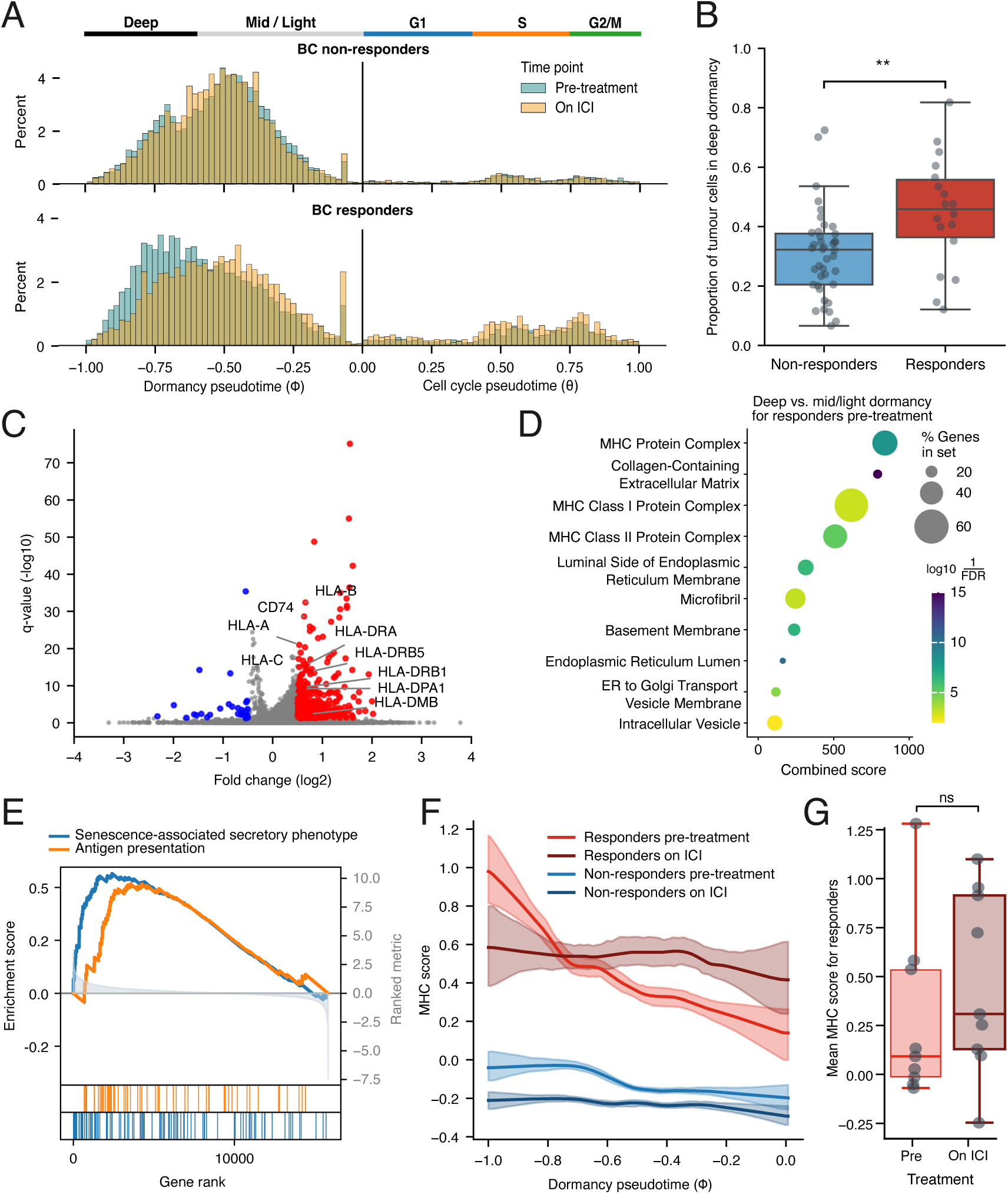
Ouroboros reveals preferential immune targeting of cells in deep dormancy for breast cancer patients treated with immune checkpoint inhibitors. (**A**) Tumour cell frequency distributions along pseudotime pre- and on-treatment for non-responders (top, n=20 patients) and responders (bottom, n=9 patients). Following treatment, responders exhibit a shift in tumour cell pseudotime from deep dormancy to mid/light dormancy. (**B**) Breast cancer responders show a higher proportion of tumour cells in deep dormancy (*ϕ <* −0.6) than non-responders. (**C**, **D**, **E**) volcano plot, top ten GO terms, and GSEA analysis show enrichment of antigen processing and presentation in deep dormancy vs. mid and light dormancy for BC responders in the pre-treatment condition. GSEA also shows enrichment of a senescence-associated secretory pheontype gene set. (**F**) Local regression for MHC score across dormancy pseudotime shows a significant negative correlation in responders pre-treatment (Spearman’s *ρ* =−0.19, p = 6.57 × 10*^−^*^63^). MHC score for cells in deepest dormancy decreases on ICI. Shading represents 95% confidence interval from 100 bootstrap resamples. (**G**) Without accounting for dormancy pseudotime, there is no significant difference in MHC score for responders pre- and on-treatment.

To examine what may be driving the loss of deep-dormant cells during ICI treatment, we identified differentially expressed genes between tumour cells in deep versus mid/light dormancy for responder samples pre-treatment. This analysis revealed that expression of major histocompatibility complex (MHC) and human leukocyte antigen (HLA) genes was elevated in deep dormancy (log_2_FC *>* 0.5, p *<* 0.05; **Figure 4C**). Gene Ontology (GO) analysis showed over-representation of terms associated with endogenous antigen processing and presentation, including the endoplasmic reticulum, vesicle transport and MHC class I and II protein complexes (log_2_FC *>* 0.5, p *<* 0.05; **Figure 4D**). Next, we asked whether elevated antigen presentation for responder tumour cells in deep dormancy was linked to a senescent phenotype. A gene set enrichment analysis (GSEA) showed gene sets associated with antigen processing and presentation (normalized enrichment score (NES) = 1.615, P = 4.944 × 10*^−^*^3^, FDR = 3.612 × 10*^−^*^3^), and the senescence-associated secretory phenotype (SASP) program (NES = 1.931, P = 0.00, FDR = 0.00) were both strongly enriched in deep dormancy (**Figure 4E**).

To determine whether antigen presentation decreases monotonically with dormancy pseudotime, we next scored MHC activity in each cell by taking the normalized average expression of canonical MHC class I (HLA-A, HLA-B, HLA-C) and class II (HLA-DRA, HLA-DPA1, HLA-DQA1) genes. MHC scores were low across dormancy pseudotime in non-responders, regardless of treatment condition (**Figure 4F**). In responder samples, MHC scores were significantly negatively correlated with dormancy pseudotime pre-treatment (Spearman’s *ρ* = −0.19, p = 6.57 × 10*^−^*^63^; **Figure 4F**). This association disappeared in responders following ICI treatment, suggesting that deep dormant cells with high MHC expression were actively eliminated (**Figure 4F**). Notably, without accounting for dormancy pseudotime, no significant change in mean MHC score with treatment was observed for responders (**Figure 4G**, **Figure S11**).

### Platinum therapy targets ovarian cancer cells in light dormancy while exposing deep dormant cells to immune targeting

To determine whether shifts in dormancy states occurred in response to other anti-cancer therapies, we next applied our model to a 10x Genomics Chromium scRNA-seq dataset derived from a cohort of advanced high grade serous ovarian cancer patients treated with chemotherapy (HGSOC, n = 11 patients, 51786 cells^39^). Zhang et al. profiled the primary tumour biopsy (“treatment-naive” time point), as well as the surgically resected tumour following three cycles of platinum-taxane neoadjuvant chemotherapy (NACT, “post-NACT” time point)^39^. Patients, cell types, and treatment time points were well-integrated in Ouroboros latent space (**Figure S12**). We first compared dormancy pseudotime distributions for the three major cell subsets annotated by Zhang et al. (**Figure 5A**). Stromal cells overwhelmingly mapped to Ouroboros deep dormancy pseudotime (median *ϕ* = −0.74) and showed little to no proliferation, consistent with terminal differentiation or senescence in this older patient cohort (median age 67 years). The tumour and immune populations, on the other hand, had a minor cycling cell subset and peaked in mid-dormancy (median *ϕ* = −0.50 and *ϕ* = −0.47, respectively), with substantial variance across the dormancy pseudotime range (immune cell 10-90th quantiles: −0.65 to −0.27, tumour cell 10-90 quantiles: −0.68 to −0.35; **Figure 5A**).

**Figure 5.**
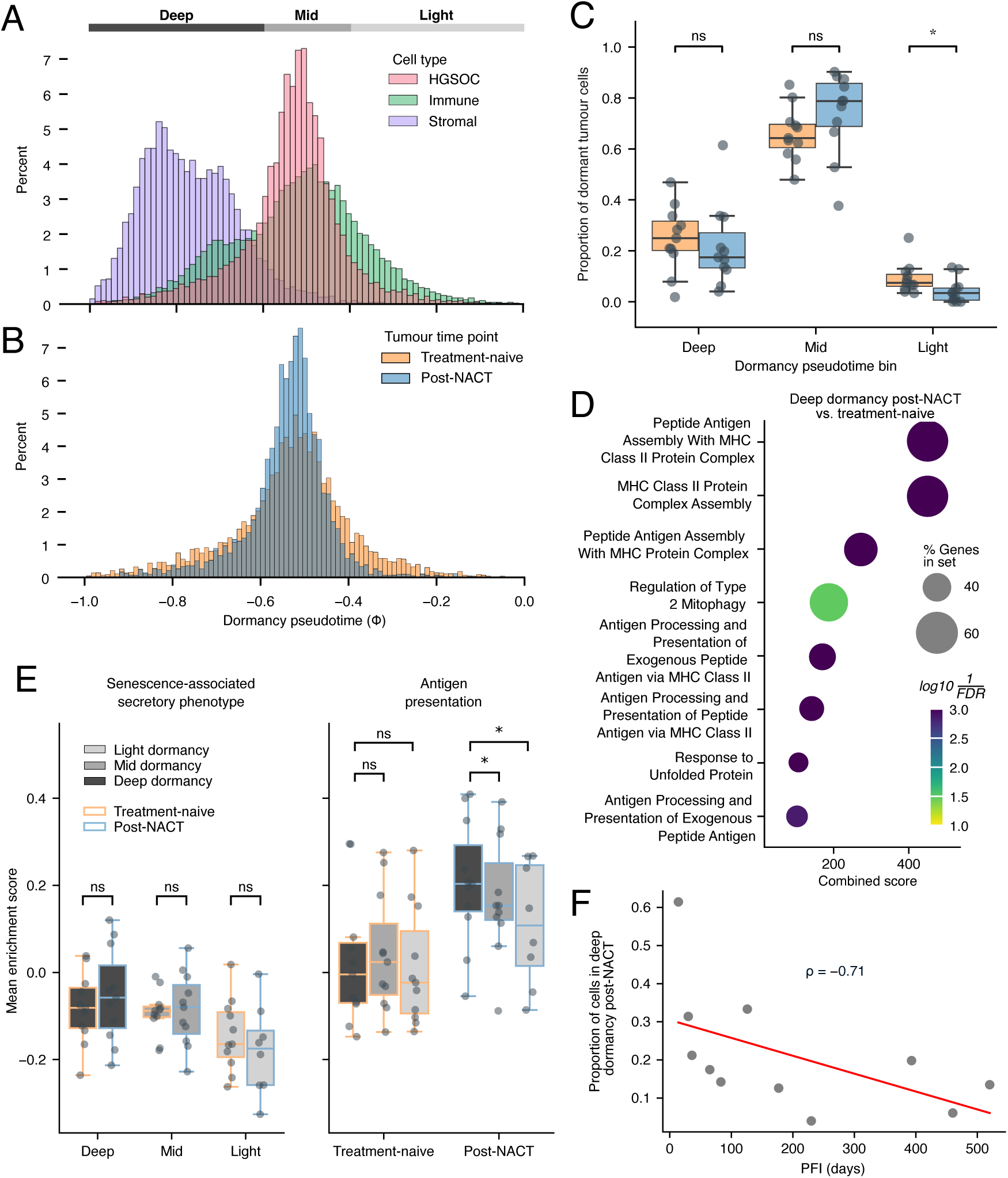
Ouroborous reveals high grade serous ovarian tumour cells converge towards mid-dormancy post-NACT treatment. (**A**) Cell frequency distribution over dormancy pseudotime for the HGSOC dataset (n=11 patients), coloured by cell type. Stromal cells primarily map to deep dormancy, while the tumour and immune distributions span from light to deep dormancy, with a peak in mid-dormancy. (**B**) HGSOC tumour cell distribution over dormancy pseudotime, coloured by treatment time point, showing a depletion of cells in both light and deep dormancy post-NACT. (**C**) Across patients, the proportion of tumour cells in light dormancy decreases significantly post-NACT (Wilcoxon test, *p* = 0.013). (**D**) GO analysis of upregulated genes (log_2_FC *>* 0.5, *p <* 0.05) for tumour cells in deep dormancy post-NACT vs. pre-treatment reveals enrichment for terms related to antigen presentation. (**E**) GSVA SASP scores increase with deeper dormancy regardless of treatment timepoint. Antigen presentation scores do not differ across dormancy depth pre-treatment, but are both elevated and associated with dormancy depth post-NACT. (**F**) A higher proportion of tumour cells in deep dormancy post-treatment is associated with shorter progression free interval (Spearman’s *ρ* = −0.71, *p* = 0.01).

Comparing the proportion of cycling tumour cells before and after treatment, we observed a substantial decrease in cell proliferation post-NACT, consistent with the primary mode of action for platinum-taxane therapeutics (**Figure S13**).

We next compared dormancy pseudotime distributions for the tumour cell population across treatment time points. Contrary to expectations, we did not observe a unidirectional shift towards deeper dormancy with chemotherapy treatment. Instead, we found a bidirectional decrease in variance post-NACT, with depletion of tumour cells at both dormancy extremes relative to treatment-naive cells (Levene’s test F = 650.84, p = 2.69 × 10*^−^*^138^; **Figure 5B**). The decrease in the proportion of cells within the light dormancy range was significant at the sample level, consistent with the primary mode of action of chemotherapy in targeting frequently cycling cells (*ϕ* = −0.4 to 0; Wilcoxon rank-sum test: W = 6.0, p = 0.0137; **Figure 5C**). The majority of patients showed a decrease in the proportion of deep dormant cells with treatment (n=7/11 patients), but response was heterogeneous and not statistically significant at the cohort level (*ϕ* = −1 to −0.6; Wilcoxon rank-sum test: W = 22.0, p = 0.365; **Figure 5C**). Given the modest size and varying response to treatment in this primary patient dataset, we sought to identify a driver for this potential distinct mode of therapeutic action.

Informed by the immunotherapy-treated breast cancer analysis (**Figure 4**), we hypothesized that chemotherapy directly targets cycling ovarian cancer cells and those in light dormancy, while inducing an immune response against cells in deep dormancy. Chemotherapy-induced immunogenic cell death (ICD) has been associated with the release of damage-associated molecular patterns (DAMPs) and immunostimulation^40^. We performed differential gene expression analysis between cells in deep and mid-dormancy post-NACT, as well as between cells in deep dormancy pre- and post-NACT, and found increased expression of genes involved in antigen processing and presentation for both comparisons (log_2_FC *>* 0.5, *p <* 0.05; **Figure S14**). In line with expectations, GO term enrichment analysis for genes upregulated in deep dormancy for the post-NACT time point relative to the pre-treatment time point was once again dominated by terms associated with antigen processing and presentation (log_2_FC *>* 0.5, *p <* 0.05; **Figure 5D**).

To comprehensively assess antigen presentation across dormancy states (deep, mid, and light) and treatment time points, we performed gene set variation analysis (GSVA) using the antigen processing and presentation and SASP gene sets. SASP enrichment scores increased modestly with deeper tumour cell dormancy, but there was no significant difference in SASP scores pre-treatment versus post-NACT (Wilcoxon rank-sum test: light dormancy, W = 8.0, p = 0.195; mid dormancy, W = 32.0, p = 0.966; deep dormancy, W = 32.0, p = 0.966; **Figure 5E**). On the other hand, antigen presentation scores showed no association with dormancy depth pre-treatment (deep vs mid: W = 31.0, p = 0.898; deep vs light: W = 18.0, p = 0.206), but were both significantly elevated and correlated with dormancy depth post-NACT (deep vs mid: W = 9.0, p = 0.0322; deep vs light: W = 2.0, p = 0.0234; **Figure 5E**). This indicates that platinum-taxane chemotherapy preferentially upregulates antigen presentation programs for tumour cells mapping deeper along dormancy pseudotime, in a manner that is distinct from general senescence-associated programs. Finally, we asked whether the efficacy of immune targeting of cancer cells in deep dormancy is associated with patient outcomes. The proportion of tumour cells in deep dormancy for each sample post-NACT sample showed a significant negative correlation with progression-free interval (PFI), suggesting that failure to launch a chemotherapy-induced immune response against deep dormant tumour cells negatively impacts treatment outcomes (**Figure 5F**).

## Discussion

Elegant molecular and imaging studies introduced the concept of the G0 continuum, a spectrum of dormancy states whereby the likelihood of cell cycle re-entry decreases with distance from the restriction point^18, 19^. In training a probabilistic deep learning model that can project heterogeneous single cell transcriptomic profiles into a single, interpretable latent space, we enable, for the first time, large-scale quantitative analysis along the G0 continuum via a common dormancy pseudotime.

Ouroboros infers cellular location along common pseudotime trajectories capturing both the active cell cycle (*θ*) and deepening cell dormancy (*ϕ*), as well as discrete cell cycle and G0 state labels. In benchmarking experiments on a FUCCI-labelled dataset, Ouroboros out-performed published methods in inferring active cell cycle phases (G1, S, G2/M) and continuous cell cycle pseudotime (*θ*). Inferring dormancy pseudotime across diverse biological contexts (cell lines, mouse embryonic development, primary human tissues) and technologies, we validated the association between deepening dormancy pseudotime and cell cycle arrest.

Applying dormancy pseudotime analysis to two cancer patient cohorts, we characterized the distinct cell dormancy states targeted by two classes of anti-cancer therapeutics. In a breast cancer cohort treated with immune checkpoint inhibitors^38^, dormancy pseudotime analysis revealed that ICI responders (defined by T cell clonotype expansion) have a higher proportion of tumour cells in deeply arrested states than non-responders. Following ICI treatment, non-reponders showed no change in dormancy pseudotime distribution, while the responder group showed a pronounced shift towards mid and light dormancy. Antigen presentation was associated with dormancy depth in responders pre-treatment, but this association was absent post-treatment, suggesting ICI induces an immune response not simply towards cells with MHC expression, but specifically towards cells in deep dormancy with high antigen presentation. These findings support observations in recent work by Marin et al., which showed that patient-derived tumour-infiltrating lymphocytes (TILs) had stronger activation *ex vivo* when presented with autologous senescent cancer cells than autologous non-senescent cancer cells^41^. The mid-dormancy state dominating the dormancy pseudotime distribution of non-responders likely corresponds to the quiescent cancer cell (QCC) phenotype described by Baldominos et al., which used a reporter linked to p27 in mouse models to identify QCC niches characterized by immune exclusion and exhaustion in triple-negative breast cancer^7^.

In a high-grade serous ovarian cancer cohort treated with platinum-taxane chemotherapy^39^, our model revealed an unexpected bi-directional inward shift in tumour cell dormancy pseudotime post-treatment. Platinum-taxane treatment triggers cell cycle arrest and apoptosis in actively proliferating cells^42^. As such, the pronounced depletion of tumour cells within the light quiescent range of dormancy pseudotime was expected. However, we also observed a more modest depletion in the deep dormancy range. Suspecting immune involvement, we compared SASP and antigen presentation scores across dormancy pseudotime. SASP expression was correlated with dormancy depth but showed no difference between timepoints, indicating that chemotherapy treatment does not induce a senescent secretory phenotype. Antigen presentation showed no association with dormancy depth pre-treatment, but increased substantially throughout dormancy pseudotime and was correlated with dormancy depth post-NACT. That chemotherapeutics can trigger immunogenic cell death (ICD) and stimulate an immune response is well-known^40^, but to our knowledge, ours is the first study to indicate that the two modes of action of chemotherapeutics target distinct tumour cell populations – inducing direct depletion of frequently cycling cells in light dormancy, while triggering an immune response towards cells in deep dormancy. We also found that the proportion of tumour cells remaining in deep dormancy post-NACT is significantly negatively correlated with progression free interval. This suggests that the efficacy of the chemotherapy-induced immune response in clearing dormant cancer cells is important for patient outcomes.

Various past studies report that treatment with chemotherapeutics can induce antigen expression in normal and malignant cells^40, 41, 43, 44^. In fact, chemotherapeutic agents are frequently applied specifically to induce and characterize the senescent phenotype^31, 41^. Our analysis of the ovarian cancer cohort shows that, in the absence of chemotherapy, SASP expression is associated with dormancy depth but antigen presentation is not. This suggests that SASP and antigen presentation are not inherently linked, and that past studies that used chemotherapeutics to study senescence may in fact be characterizing a particular phenotype that is not representative of senescent states more generally. Dormancy pseudotime inference now permits quantitative analysis of the full spectrum of cell dormancy states without the need for senescence-inducing drugs.

We expect our model to be widely applied to identify the dormancy states targeted by other anti-cancer therapeutics. This could facilitate efforts to identify effective drug combinations that together target cells across a broad spectrum of dormancy states to prevent relapse. As the cohort sizes of single cell cancer studies increase, dormancy pseudotime analysis could also reveal whether patient-specific dormancy distributions pre-treatment effectively predict therapeutic response, and help identify the most efficacious treatment course or order of application of therapeutics for each individual. Future studies will elucidate the genetic and epigenetic underpinnings of patient tumour dormancy states, including their relationship with cancer subtype, mutation status, clonal composition, and patterns of ongoing genomic and chromosomal instability.

Given the ubiquity of single cell transcriptomic datasets and the essential role that cell cycle and dormancy dynamics play in development, tissue repair, immune function and aging, we anticipate that dormancy pseudotime inference will unlock new insights across the life sciences.

## Methods

### Data processing

FASTQ files for all datasets except the breast and ovarian cancer cohorts were downloaded from SRA using the accession number available in the original publication. The ovarian and breast cancer cohort read count matrices were downloaded from GEO and http://biokey.lambrechtslab.org, respectively.

Datasets with cell barcodes and unique molecular identifiers (UMIs) were aligned to either hg38 or m39, and count matrices were generated using STARSolo (2.7.11a). A custom workflow was used to process the U2OS (Smart-seq2) libraries to extract spliced and unspliced counts (see Code availability).

Scanpy (v1.8.2) was used for data quality control and normalization. Poor quality cells were first filtered using the criteria applied in the original publication for each dataset. Further filtering was performed to remove cells lacking expression across all Ouroboros feature genes. ScDblFinder (v.1.16.0) was applied to identify and filter doublets and SoupX (v1.6.2) was used to remove ambient RNA contamination from droplet-based libraries. Expression values were normalized with the Scanpy function `sc.pp.normalize total`, followed by log-transformation and scaling.

### Assigning phase labels for training and benchmarking

FUCCI marker gene expression was available for the U2OS^25^ and RPE1^30^ datasets. Mahdessian et al.^25^ applied thresholds to the z-score normalized red and green fluorescent protein measurements to assign discrete cell cycle phase (G1, S, G2/M) for the U2OS dataset, and we applied the same normalization and thresholds to assign discrete phases to cells in the RPE1 dataset (**Figure S6**).

To distinguish G0 cells in the FUCCI datasets and assign cell cycle phase in unlabelled datasets used for feature selection and training, we used the following process. First, we identified annotated cell cycle genes (GO accession ‘GO:0007049’) and genes associated with G0 in the literature^16, 28, 29, 33, 45, 46^, for a total of 2120 cell cycle-associated genes or their 2188 mouse orthologs (**Table S1**). We subset each dataset to this gene set and performed Leiden clustering with Scanpy. For the RPE1 dataset, Harmony^47^ (v0.0.10) was applied before clustering to correct for batch effects between experimental conditions (untreated, pulse and chase plates). ScVelo (v0.2.3) was used to estimate and plot RNA velocity vectors. **Figure S15** shows standard UMAPs based on highly variable genes along side UMAPs based on the cell cycle gene list for four datasets (U2OS^25^, fibroblast^16^, pancreatic endocrinogenesis^34^, and RPE1^30^). For datasets without FUCCI labels, Leiden clusters with high expression of well-characterized S and G2/M phase marker gene sets^28^ were labelled as S and G2/M, respectively. For all datasets, Leiden clusters between the S and G2/M phases that maintained expression of the Graham quiescence downregulated gene set^29^ and had RNA velocity consistent with progression through the cell cycle were labelled as G1 phase. Clusters adjacent to G1 with low expression of S and G2/M marker gene sets, lower expression of the Graham quiescence downregulated gene set, and RNA velocity direction consistent with cell cycle exit were labelled as ‘G1-G0 transition’. Finally, clusters with near-absent expression of cell cycle and quiescence downregulated genes and no clear direction in RNA velocity were labelled as G0 (**Figure S15**). In the RPE1 dataset we used the label ‘non-cycling’ to encompass both G1-G0 transition and G0 cells, as this dataset had low overall expression and RNA velocity direction was not sufficiently clear to distinguish these two subpopulations (**Figure S15**).

### Feature selection and model training

Feature selection was performed based on the U2OS, fibroblast and mouse pancreatic datasets. Mouse genes were first converted to their human orthologs or discarded if no orthologs were found (Ensembl version 114). We applied a Wilcoxon rank-sum test to identify genes with significantly higher expression for each phase in each dataset. Of those with significant difference in expression, the 8% of genes with largest log-fold change values were selected for each phase and dataset. Genes that were common between all three datasets were retained to form a preliminary training feature set. Next, we trained a preliminary model using scPhere^24^ with the von Mises-Fisher (vMF) distribution and a 3D latent space on the unprocessed gene expression counts for 70% of cells from the combined U2OS, pancreatic and fibroblast datasets. We mapped the remaining 30% of cells into the latent space and applied k-nearest neighbours (KNN) classification (scikit-learn v.0.24.2; k = 35) to predict the phase identity of this test set. We computed SHAP (v0.42.0) values using the `shap.KernelExplainer` function applied to the trained encoder-KNN workflow and averaged absolute SHAP values across cells in each phase in the test set to assess feature importance. We selected the top 5% of absolute SHAP values for S and G2/M, and the top 15% for G1, G1-G0 transition and G0 to create our final feature set of 226 training genes (**Table S1**). Finally, we used the unprocessed counts from the U2OS and fibroblast datasets and the SHAP feature set to fit the reference Ouroboros VAE embedding.

### Discrete phase assignment, cell cycle pseudotime, and dormancy pseudotime

To assign discrete phases to unlabelled cells mapped to Ouroboros latent space, we use the k-nearest neighbours in the reference datasets (k = 25; **Figure 1A**), and select the majority label.

To define cell cycle pseudotime (*θ*), we performed principal component analysis (PCA) on the latent 3D coordinates for reference cells in the active cell cycle (G1, S and G2/M). We defined the north-south axis as the third principal component (PC), and used the plane perpendicular to this axis that cuts through the centre of the sphere (also known as a ‘great circle’) to define the equatorial line. Next, we projected all cells in the northern hemisphere onto this equatorial plane. We set *θ* = 0 = 1 at the radial line passing through the midpoint between reference G2/M and G1 cells, and defined cell cycle pseudotime as the angle around the northern pole (**Figure 1A**,**D**), **Figure S16**). This angle, *θ*, is scaled to span from 0 to 1.

To define dormancy pseudotime (*ϕ*), we first identified a point on the surface of the spherical embedding that corresponds to deepest dormancy. We applied PCA to all G1-G0 transition and G0 cells in the reference datasets. We selected the meridian line as the ‘great circle’ defined by the first two PCs. All cells were projected onto the meridian plane. Next, we grouped cells into equal-sized bins based on their angle around this circular projection. We identified the bin with minimum cell counts as the gap between the dormancy trajectory and active cell cycle. Of the two adjacent bins, we identified the one containing the ‘G0 terminus’ as the bin with highest number of G0 cells, and defined the deepest point of dormancy pseudotime as the median coordinate for cells within this bin. Finally, *ϕ* was defined as the latitudinal distance between each cell in the southern hemisphere and the ‘G0 terminus’ (where, in this instance, the coordinate reference system is redefined such that the G0 terminus is denoted as the southern pole). The pseudotime *ϕ* was then scaled to span from 0 and −1 (**Figure S17**).

### Benchmarking discrete phase and cell cycle pseudotime

FUCCI-labelled RPE1 cells from Battich et al.^30^ (chase, pulse and control experiments) were downloaded and processed as described above. This dataset was analyzed using Ouroboros and previously published tools.

The `score genes cell cycle` function from Scanpy (v1.10.2) was applied to the RPE1 cells using default options. Prior to running Tricycle, Python anndata (v0.7.5) objects were converted to R SingleCellExperiment objects with zellkonverter (v1.4.0). Tricycle (v1.10.0) was then run with default parameters, producing six cell cycle phase labels (G1.S, S, G2, G2.M, M.G1, and NA). For our benchmarking assessment, any intermediary phases (e.g. G1.S and M.G1) were accepted as a ‘correct’ labelling if either phase matched the FUCCI phase. DeepCycle (no version number available) was run using default parameters except the `expression threshold` argument was set to 0.00001 on account of the low total transcript counts in this dataset. The hotelling filter was not applied, and MELK was chosen as the training gene since this gene was used for the human dataset analyzed in the DeepCycle publication^16^. Phases were assigned to cells between the output phase boundaries, and were considered a ‘correct’ labelling if they matched the FUCCI phase. DeepCycle output three possible M/G1 phase boundaries, and the boundary with the highest percentage of correctly classified cells was used to evaluate performance (**Figure S18**).

### Breast and ovarian cancer cohort analyses

For the breast cancer cohort^38^, non-proliferating tumour cells were divided into two dormancy pseudotime bins: deep dormancy (−1 ≤ *ϕ <* −0.6), and mid/light dormancy (−0.6 ≤ *ϕ <*0). For the ovarian cancer cohort^39^, three dormancy pseudotime bins were defined: deep (−1 ≤ *ϕ <* −0.6), mid (−0.6 ≤ *ϕ <* −0.4), and light dormancy (−0.4 ≤ *ϕ <*0).

Differential gene expression analysis was performed using the `rank genes groups` function from Scanpy (v1.10.3) with the Wilcoxon method, a minimum absolute log_2_ fold-change threshold of 0.5 and a significance cut off of p = 0.05. Genes expressed in 30 or fewer cells were excluded. The GSEApy package (v1.1.9) was used to perform pathway, gene set enrichment (GSEA), and gene set variation analyses (GSVA). Gene sets were obtained from the Molecular Signatures Database (MSigDB) v2025. For the breast cancer cohort, the top ten GO Cellular Component (2025) terms comparing deep vs. mid/light dormancy for responder tumour cells pre-treatment were plotted. For the ovarian cancer cohort, the top eight GO Molecular Function terms between deep dormancy post-NACT vs. pre-treatment for tumour cells were plotted.

GSEA KEGG gene set M16004 was used to assess enrichment in antigen processing and presentation. To assess enrichment specifically for senescence-associated secretory phenotype (SASP), rather than dormancy more broadly, we filtered the SenMayo^32^ gene set (GSEA accession M45803) to retain only secreted genes (**Table S1**). Genes found in the Ouroboros feature set were excluded from pathway enrichment analysis, in order to identify enriched processes independent of dormancy pseudotime.

For the breast cohort, MHC expression scores were obtained with the Scanpy `score genes` function, using canonical genes that encode for MHC class I (HLA-A, HLA-B, HLA-C) and II (HLA-DRA, HLA-DPA1, HLA-DQA1). T cells with cell cycle pseudotime greater than 0.4, corresponding to cells in S and G2/M phases of the cell cycle, were classified as actively proliferating, in line with the S and G2/M phase scoring from the original breast cancer cohort publication^38^.

For the ovarian cohort, GSVA analysis was performed on six clusters of tumour cells, representing the deep, mid, and light dormancy pseudotime bins before (treatment-naive) and after (post-NACT) treatment.

### Sphere visualizations

Three-dimensional sphere visualizations were created using Plotly (v5.24.1). To create 2D Robinson projections of the spherical latent space, we converted the cartesian coordinates of each cell to latitude and longitude and used Cartopy (v0.23.0) to create the projection, centred at a longitude of 80*^◦^*.

### RNA velocity in spherical latent space

To generate RNA velocity vectors for a dataset in the spherical latent space, we used the jointly normalized, log-transformed and scaled gene expression values, subset to include only genes in the Ouroboros feature set. Next, we set the UMAP space in the AnnData (v0.7.5) object for the dataset to the Cartesian coordinates (x, y, z) of cells within the spherical latent space, and ran ScVelo with default parameters on this object. To generate RNA velocity vectors on the Robinson projections, we set the UMAP space in the AnnData object to the converted Robinson longitude and latitude values for each cell, as described above.

## Data availability

Data for the U2OS, fibroblast, and RPE1 datasets are available from the Gene Expression Omnibus (GEO) under accession codes GSE146773, GSE167609 and GSE128365, respectively. Processed count matrices for the BC and HGSOC patient datasets are available from http://biokey.lambrechtslab.org and GEO under accession code GSE165897, respectively.

## Code availability

The Ouroboros Python package and pre-trained computational model will be available on GitHub at https://github.com/steiflab/Ouroboros.

## Acknowledgements

We gratefully acknowledge funding support from the Canadian Institutes of Health Research (PJF185713) and BC Cancer Foundation (F202201). A.S. is a Michael Smith Health Research BC Scholar (SCH20222625).

## Author contributions

H.M. developed the feature set, trained and benchmarked the model. H.M., M.T. and G.C. performed data analyses. H.M. and G.C. developed the Python package. A.S. conceived of and supervised the project. H.M. and A.S. wrote the manuscript, with contributions from M.T. and G.C. All authors approved the final manuscript.

