## Supplementary Figures for "Deep learning inference of universal dormancy pseudotime reveals the cellular targets of anti-cancer therapies"

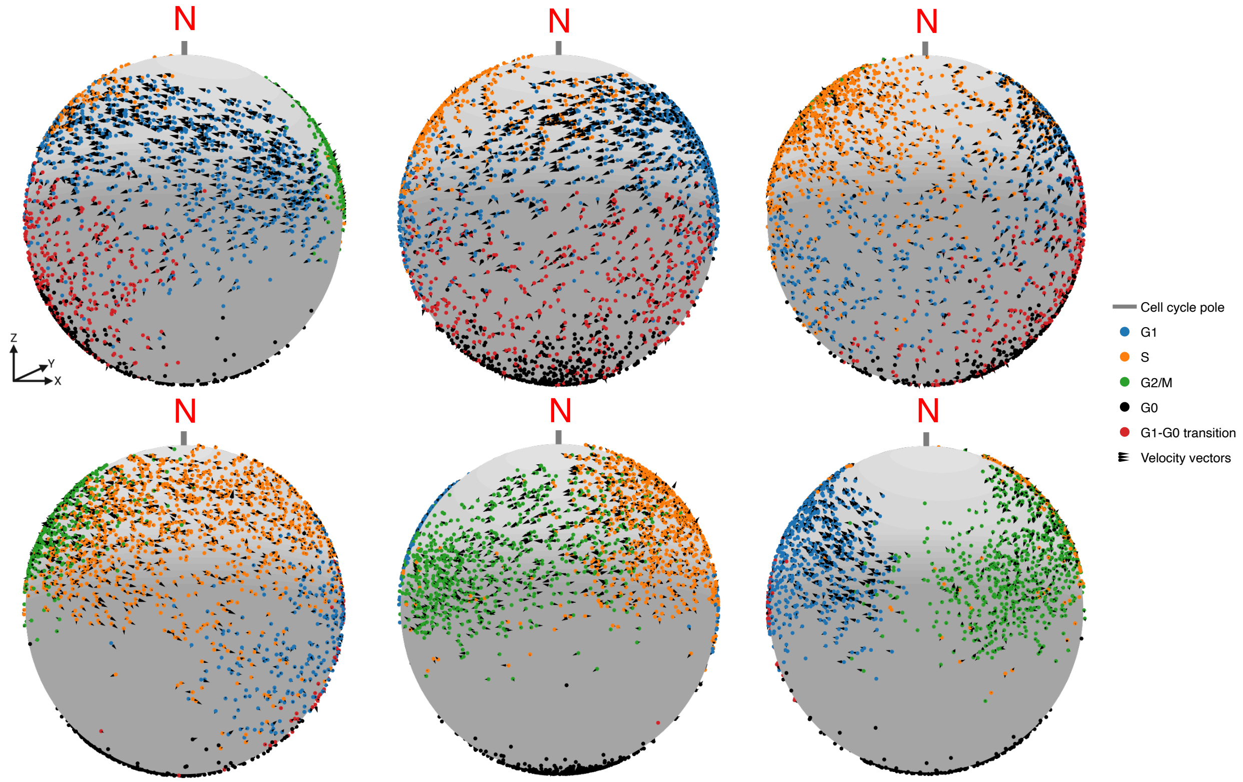


**Figure S1.** Full rotation around the Ouroboros spherical latent space with reference datasets showing cell cycle progression and RNA velocity.

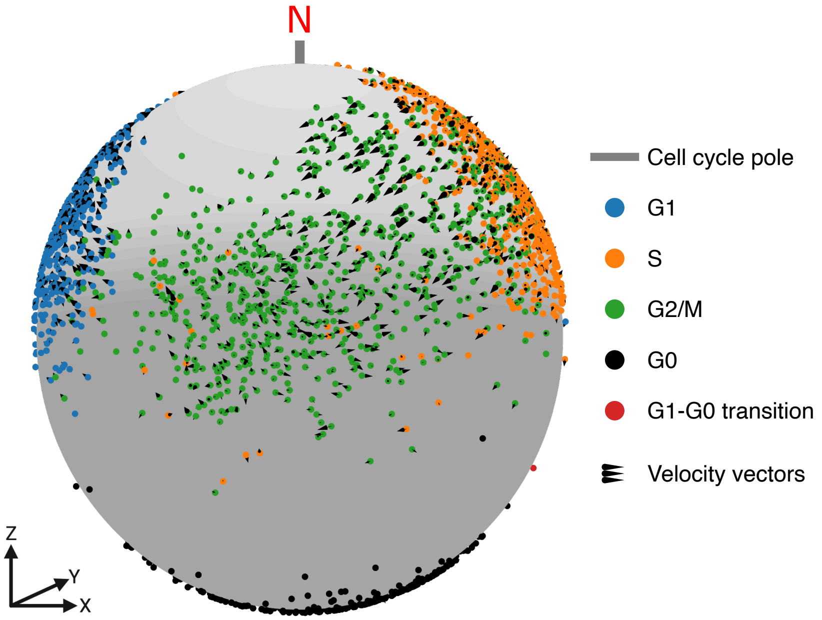


**Figure S2.** Reference dataset cell cycle embedding in the Ouroboros spherical latent space. G2/M cells have velocity vectors with low magnitude and discernible direction, consistent with the known pause in transcription at this phase. A clear gap between G2/M and G1 where is also apparent, likely corresponding to telophase or cytokinesis.

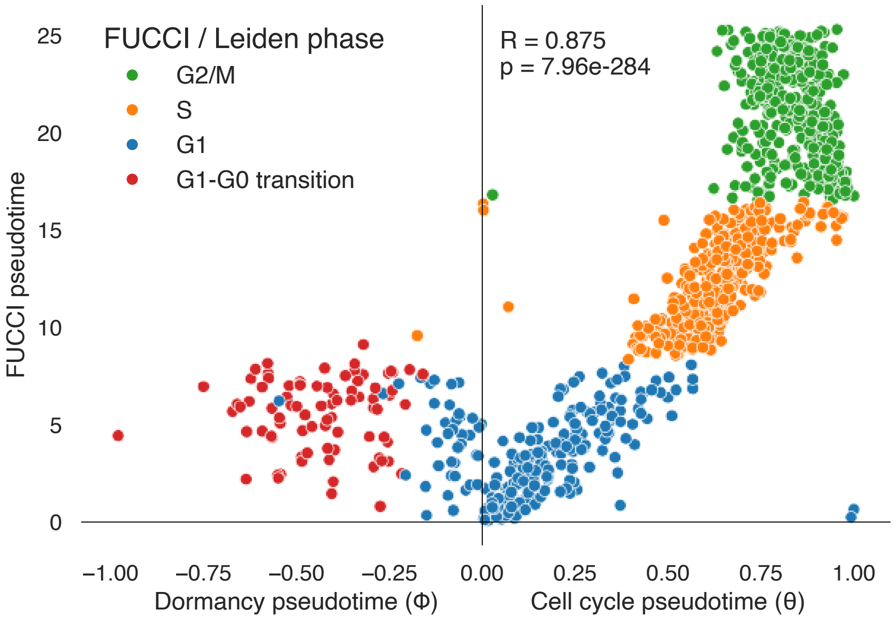


**Figure S3.** Ouroboros cell cycle and dormancy pseudotimes compared to FUCCI ground-truth cell cycle pseudotime for the U2OS training dataset. Pearson's correlation between Ouroboros cell cycle pseudotime and FUCCI cell cycle pseudotime is shown.

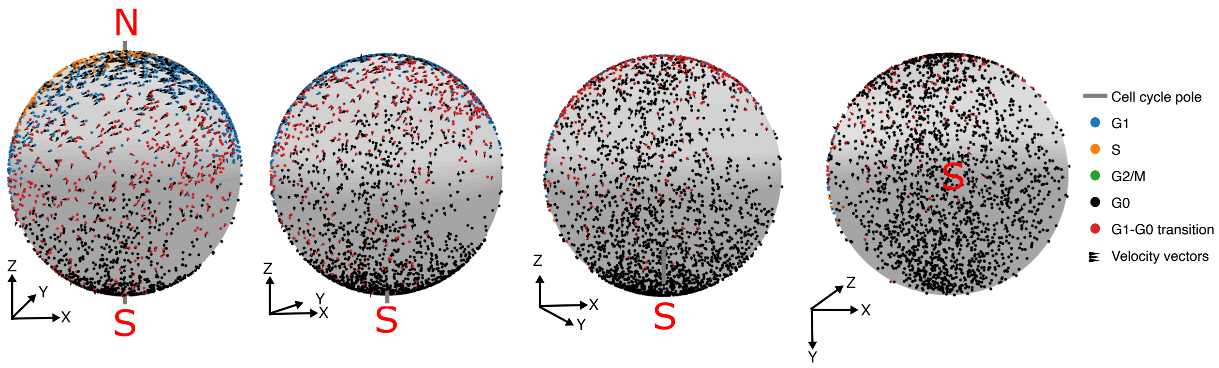


**Figure S4.** Rotation along the meridian line towards the southern hemisphere of Ouroboros latent space, showing first cells in G1-G0 transition and later in G0 extend down past the southern pole until they reach the "G0 terminus".

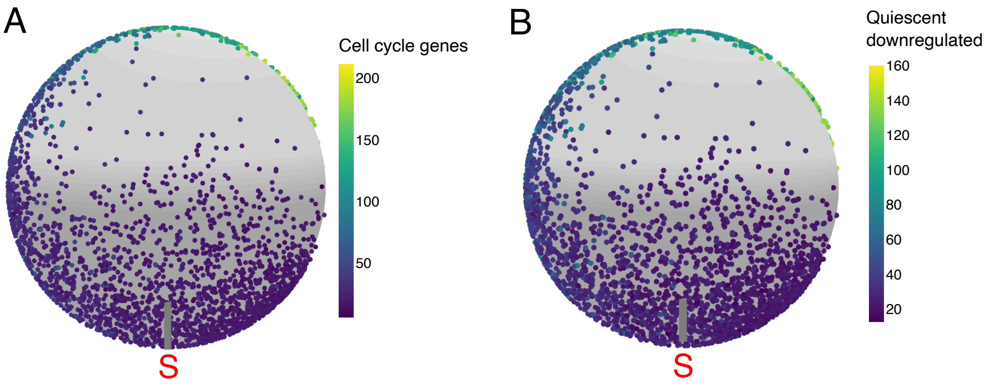


**Figure S5.** Expression of well-characterized cell cycle (A) and quiescence-downregulated genes (B) decrease with Ouroboros dormancy pseudotime.

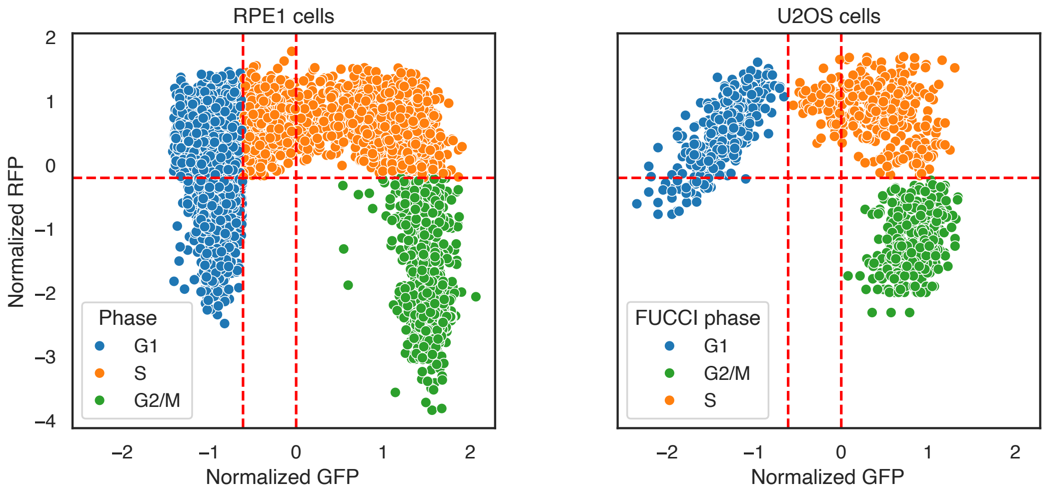


**Figure S6.** Discrete cell cycle phase labels were assigned to RPE1 cells (left) based on FUCCI marker gene fluorescence. The RFP and GFP measurements were z-score normalized, the same thresholds used by Mahdessian et al. for U2OS cells (right) were applied to both datasets.

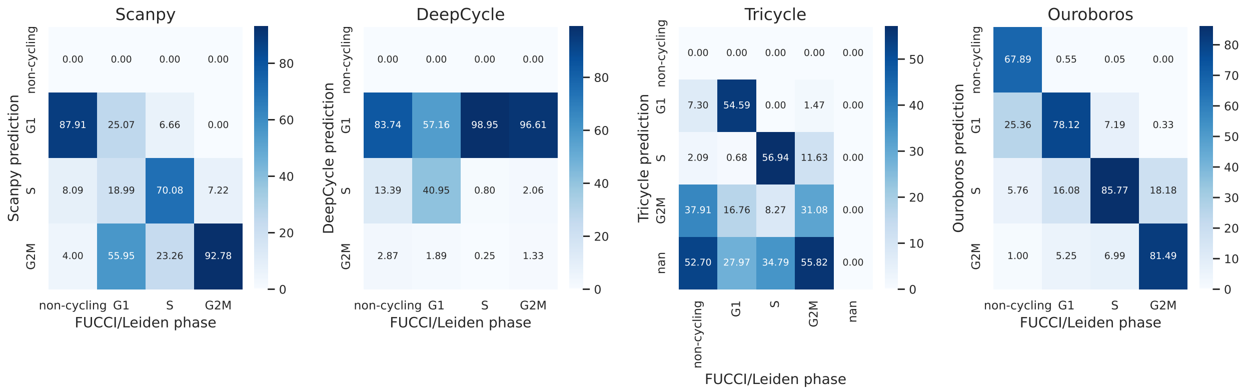


**Figure S7.** Benchmarking confusion matrices showing the performance of Scanpy, DeepCycle, Tricycle and Ouroboros on the FUCCI-labelled RPE1 dataset.

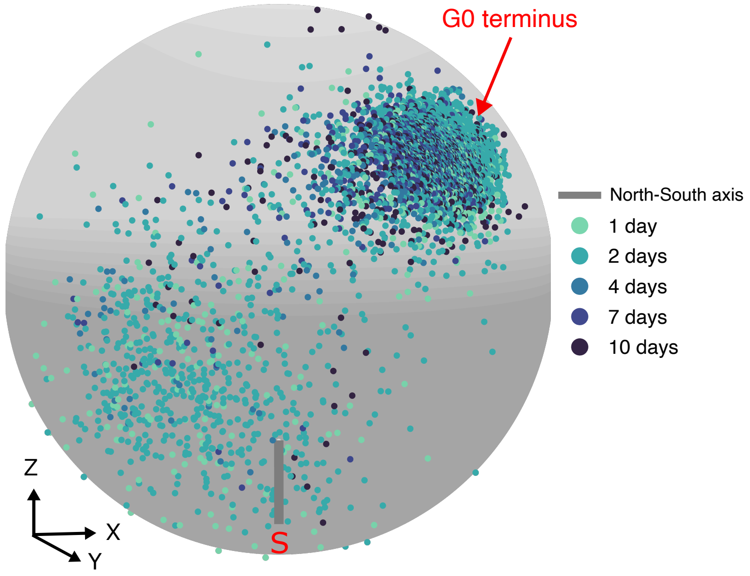


**Figure S8.** Etoposide-treated fibroblasts over a 10-day time series experiment from Wechter et al. embedded in Ouroboros latent space.

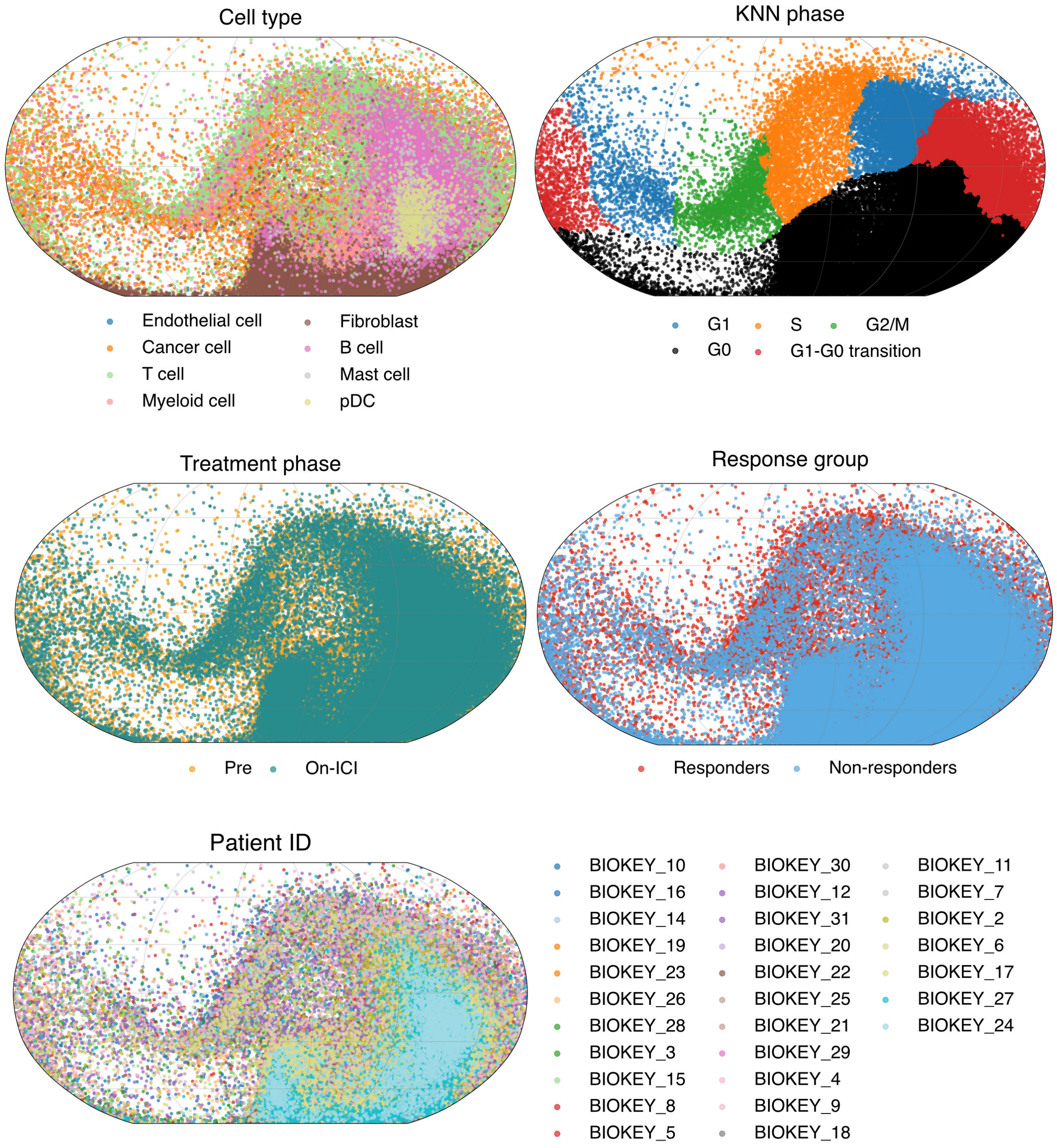


**Figure S9.** The breast cancer dataset in Ouroboros latent space, coloured by cell type, KNN label, treatment phase, response group, and patient.

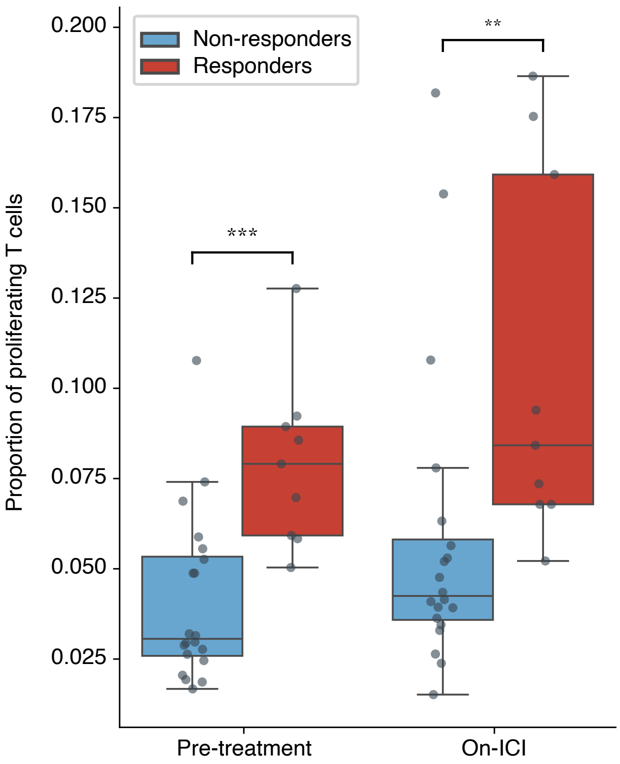


**Figure S10.** The proportion of proliferating T cells was significantly higher in the responder group than the non-responder group of the breast cancer cohort at both time points, indicating a higher level of immune activation even before initiation of ICI treatment.

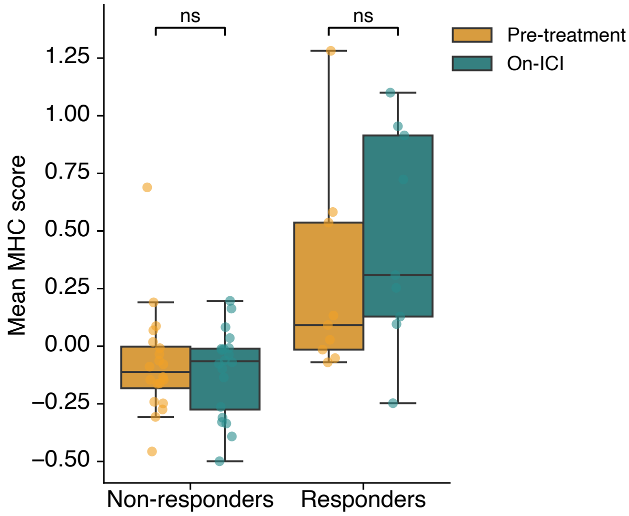


**Figure S11.** Without accounting for dormancy pseudotime, there were no significant differences in MHC scores between treatment time points in the breast cancer cohort for either the responder or non-responder group.

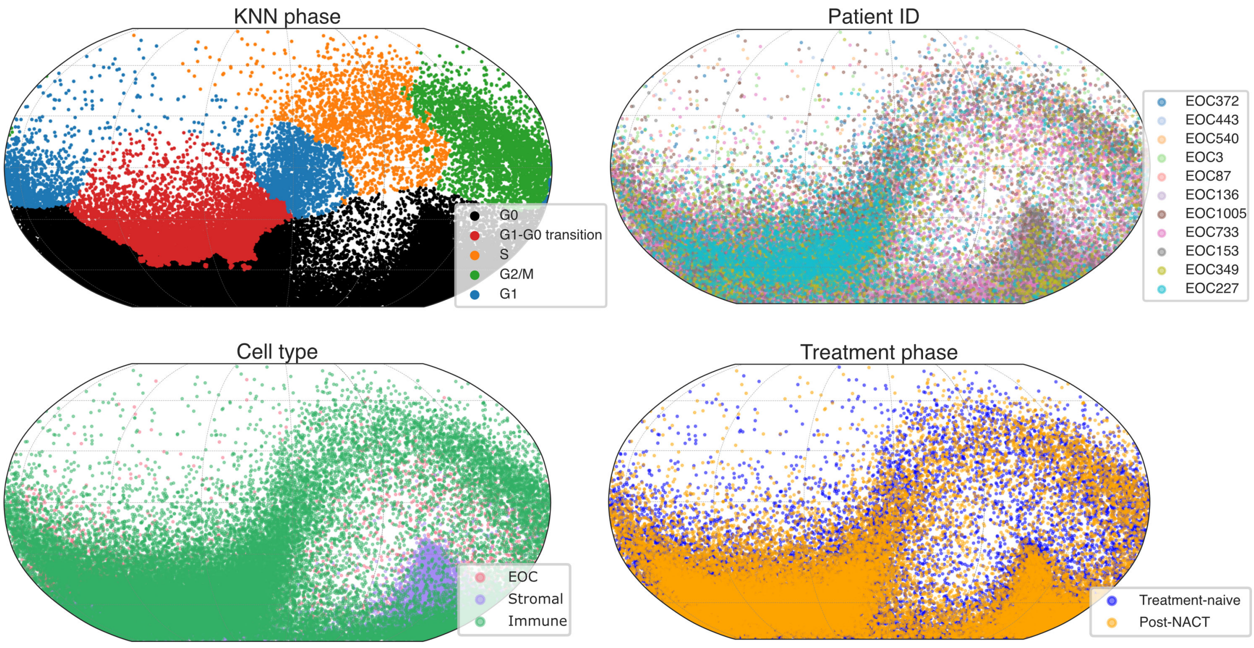


**Figure S12.** The HGSOC dataset in Ouroboros latent space, coloured by KNN label, patient, cell type, and treatment time point.

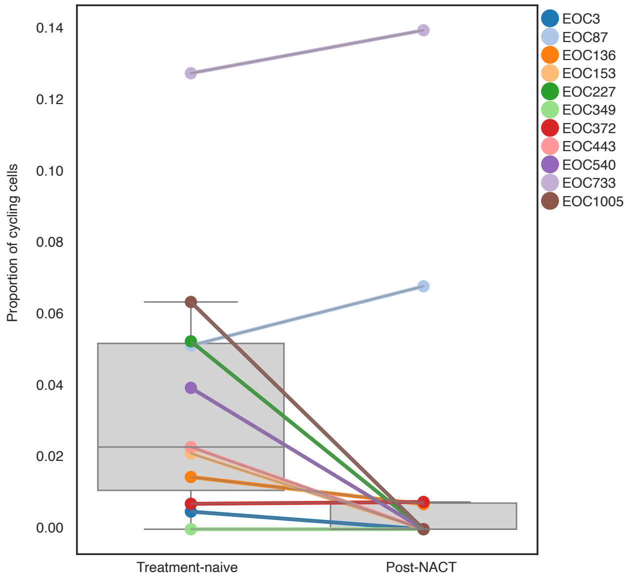


**Figure S13.** Proportion of HGSOC tumour cells in active phases of the cell cycle out of all tumour cells for each patient across treatment time points. Post-NACT, only two patients did not show a reduction in cycling cell proportion.

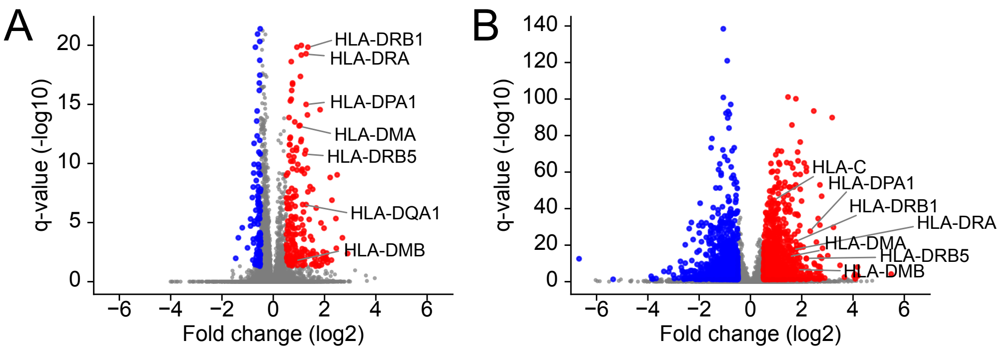


**Figure S14.** Volcano plots for HGSOC differential expression analysis. (a) Comparison between deep and mid-dormancy for the post-treatment time point. (b) Comparison between deep dormancy pre- and post-treatment.

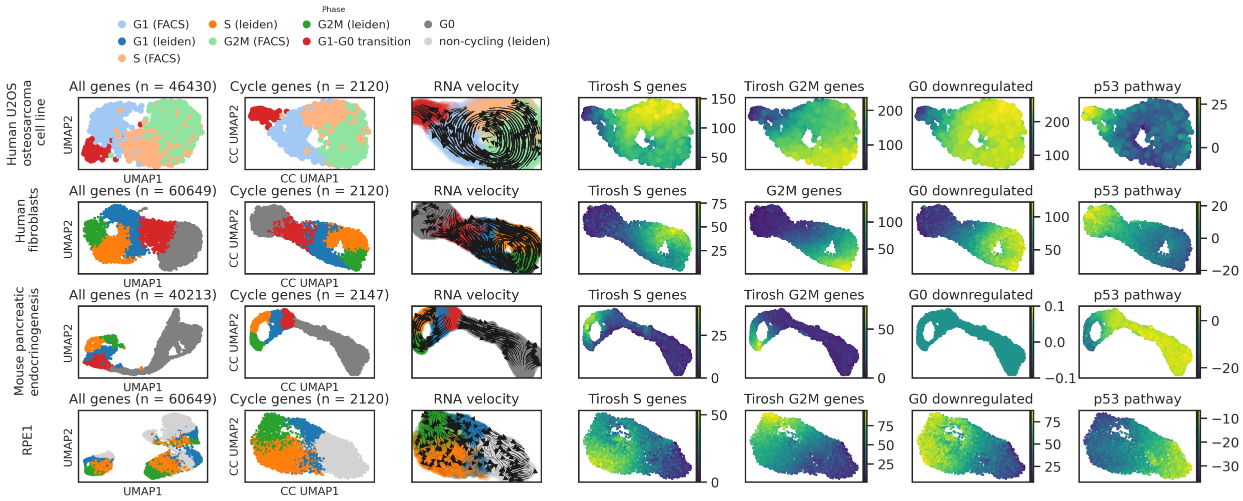


**Figure S15.** A detailed characterization of the datasets labelled with our phase assignment paradigm. The first three datasets were used for SHAP feature selection (U2OS, fibroblast, pancreatic endocrinogenesis) while the RPE1 dataset was used for testing. FUCCI labels for G1, S, and G2/M phases were available for the U2OS and RPE1 datasets. UMAPs based on cell cycle genes, RNA velocity direction, cell specific expression of the Tirosh et al. S-phase and G2M-phase gene sets, expression of the Graham quiescent down gene set, and GSEA enrichment score for the p53 pathway. Differences in the number of genes used for UMAP projection between the datasets result from differences in the mouse and human genomes and their respective annotations.

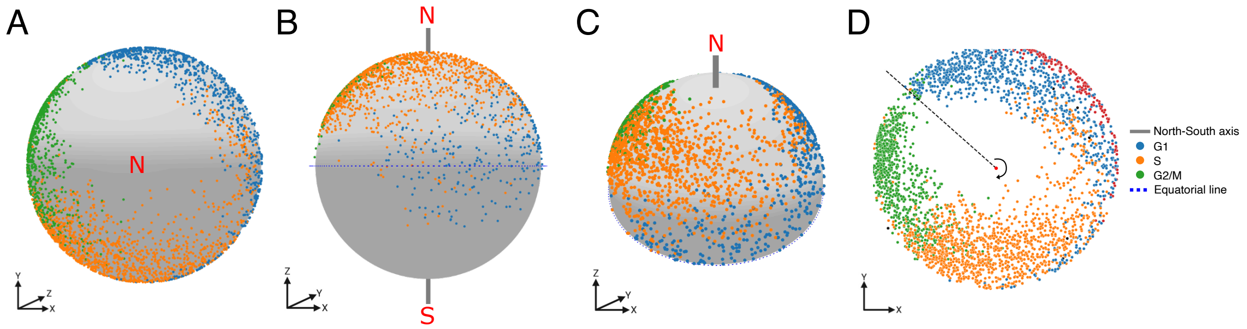


**Figure S16.** Identifying the equatorial line, North-South axis, and cell cycle pseudotime in Ouroboros latent space. (a) View of the reference embeddings in Ouroboros latent space looking down the north pole. (b, c) To identify the equatorial plane, PCA was applied to all cells labelled G1, S and G2/M in the training datasets based on their coordinates, and the third PC was taken as the equatorial plane. The northern pole was selected as the centroid of these cycling cells. (d) All points in the northern hemisphere were projected down onto the equatorial plane to calculate cell cycle pseudotime.

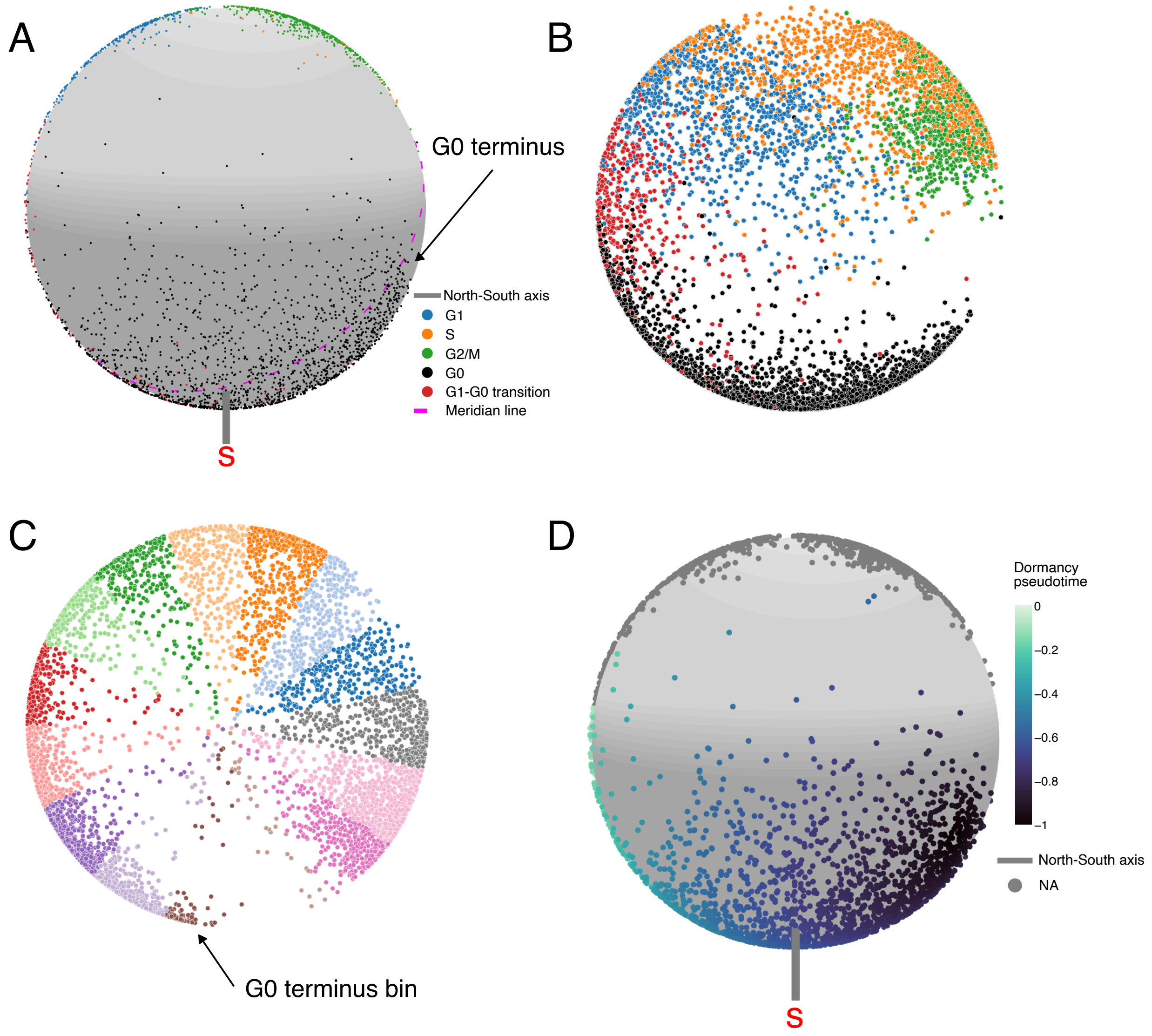


**Figure S17.** Identifying the meridian line and dormancy pseudotime in Ouroboros latent space. (a, b) Demonstration of PCA-based meridian line drawn through the majority of non-cycling cells and the projection of all cells onto this plane. (c) The cells were binned by angle around the plane, and the G0 terminus was determined by identifying the bin with fewest cells in the reference dataset, and then selecting the median spherical 3D coordinates of cells in the preceding bin as the terminus. (d) Dormancy pseudotime was determined based on the normalized longitudinal distance from the G0 terminus, and was assigned only to cells in the southern hemisphere.

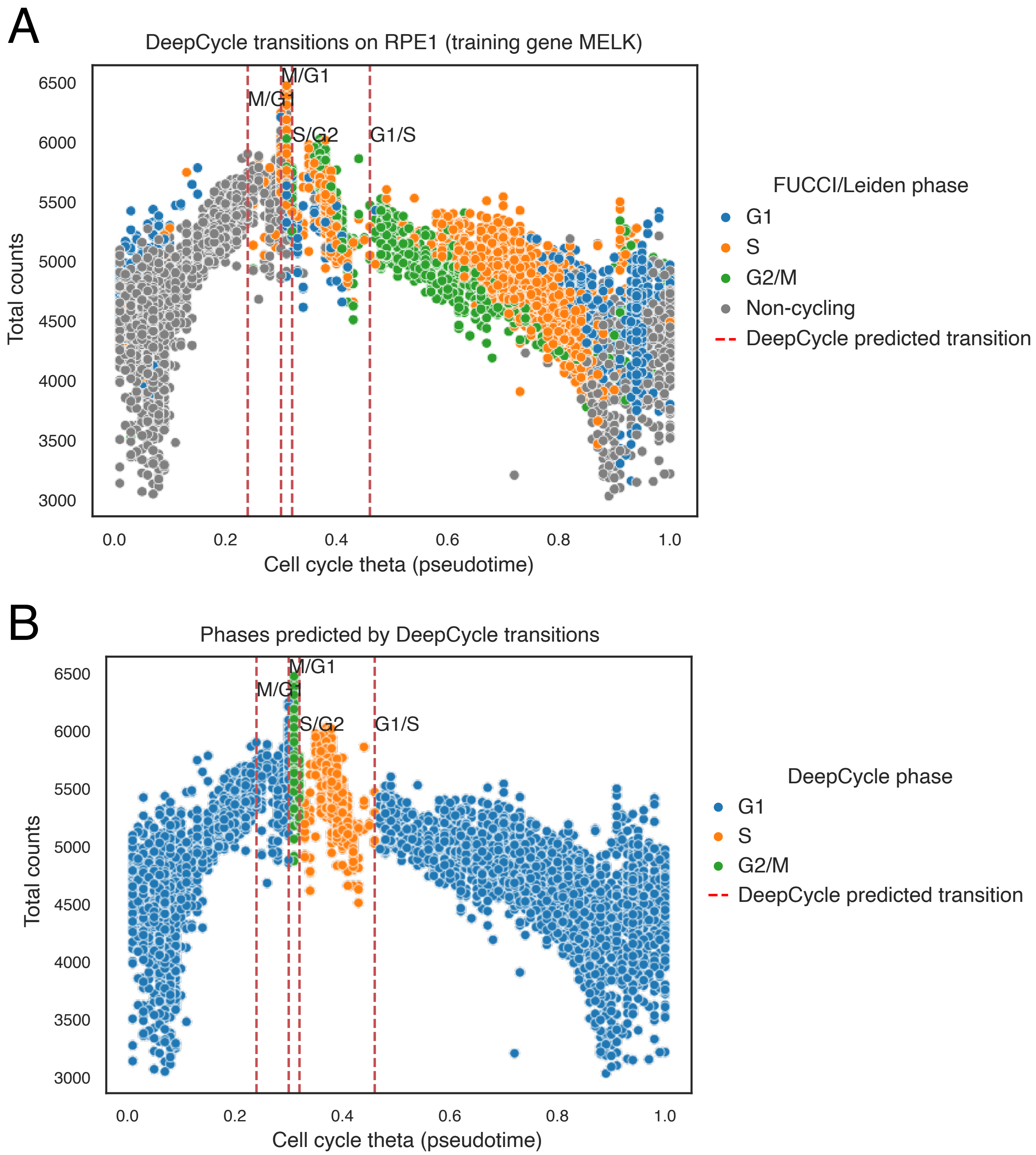


**Figure S18.** DeepCycle results on the RPE1 FUCCI-labelled dataset. (a) DeepCycle cell cycle pseudotime trajectory, coloured by FUCCI / Leiden phase labels. Dotted red lines represent the phase transitions predicted by DeepCycle, which are used to assign class labels. (b) The DeepCycle trajectory coloured by DeepCycle assigned phase labels based on the phase transition boundaries.
